# Biochemical activity is the default DNA state in eukaryotes

**DOI:** 10.1101/2022.12.16.520785

**Authors:** Ishika Luthra, Xinyi E. Chen, Cassandra Jensen, Abdul Muntakim Rafi, Asfar Lathif Salaudeen, Carl G. de Boer

## Abstract

Genomes encode for genes and the regulatory signals that enable those genes to be transcribed, and are continually shaped by evolution. Genomes, including those of human and yeast, encode for numerous regulatory elements and transcripts that have limited evidence of conservation or function. Here, we sought to create a genomic null hypothesis by quantifying the gene regulatory activity of evolutionarily naïve DNA, using RNA-seq of evolutionarily distant DNA expressed in yeast and computational predictions of random DNA activity in human cells and tissues. In yeast, we found that >99% of bases in naïve DNA expressed as part of one or more transcripts. Naïve transcripts are sometimes spliced, and are similar to evolved transcripts in length and expression distribution, indicating that stable expression and/or splicing are insufficient to indicate adaptation. However, naïve transcripts do not achieve the extreme high expression levels as achieved by evolved genes, and frequently overlap with antisense transcription, suggesting that selection has shaped the yeast transcriptome to achieve high expression and coherent gene structures. In humans, we found that, while random DNA is predicted to have minimal activity, dinucleotide content-matched randomized DNA is predicted to have much of the regulatory activity of evolved sequences, including active chromatin marks at between half (DNase I and H3K4me3) and 1/16th (H3K27ac and H3K4me1) the rate of evolved DNA, and the repression-associated H3K27me3 at about twice the rate of evolved DNA. Naïve human DNA is predicted to be more cell type-specific than evolved DNA and is predicted to generate co-occurring chromatin marks, indicating that these are not reliable indicators of selection. However, extreme high activity is rarely achieved by naïve DNA, consistent with these arising via selection. Our results indicate that evolving regulatory activity from naïve DNA is comparatively easy in both yeast and humans, and we expect to see many biochemically active and cell type-specific DNA sequences in the absence of selection. Such naïve biochemically active sequences have the potential to evolve a function or, if sufficiently detrimental, selection may act to repress them.

## Introduction

The genome encodes for many biochemical activities that can be reliably measured in cells. While some of these activities are functional and critical to organismal fitness (e.g. transcription of an essential gene), others may not be. Long non-coding RNAs (lncRNAs) are a class of RNA that are often spliced and expressed at low levels and in tissue-specific ways (1–5). While some lncRNAs are known to have critical functions, many more lncRNAs exist and debate is ongoing about what fraction of these are functional (2,6–13). More generally, 80% or more of the genome produces reproducible chromatin marks (e.g. histone modifications) (14), but genetic data suggests that closer to 10% of the genome is functional in the sense that its sequence impacts organismal fitness (15,16), and much of the pervasive transcription of genomes is consistent with noise inherent to the transcriptional machinery (7,17). Eddy proposed a “Random Genome Project” thought experiment as a means to calibrate our expectations for what activities we expect a genome to display in the absence of selection, suggesting that exogenous and evolutionarily distant or random DNA could be used as a means for generating this null (18).

There has since accumulated a great deal of evidence that random DNA can have gene regulatory activity. Certain *in vitro* transcription factor (**TF**) specificity-characterization assays rely on the ability of TFs to find their cognate site in a pool of random sequences (19,20). Random sequences have also been shown in reporter assays to have *cis*-regulatory activity in bacteria (21), yeast (22,23), human (24,25), and *Drosophila* (26), as well as 5’UTR (27,28), polyadenylation (29), and splicing activities (30), suggesting that gene regulatory sequence signals can appear frequently by chance. Yeast transcripts have been predicted to occur frequently in random DNA (31), and shown to emerge in semi-random DNA at ∼10-200 kb scales (31,32). However, no one has created a proper “null hypothesis” that captures the frequency at which we expect to see gene regulatory structures and biochemical activities in the absence of selection in yeast or humans.

Here, we create a draft null hypothesis for DNA regulatory activity in the absence of selection by profiling the gene regulatory activities of naïve and evolved DNA in both yeast and humans (**Fig. 1**). Surprisingly, evolutionarily distant exogenous DNA expressed in yeast (*Saccharomyces cerevisiae*) encodes for a plethora of transcripts covering essentially the entire DNA sequence. The naïve transcripts have similar lengths and expression levels to the evolved yeast genes and do not reflect the regulatory functions from the source organism (human), indicating that the naïve and evolved transcripts share a biochemical origin that is encoded distinctly in yeast.

**Figure 1:**
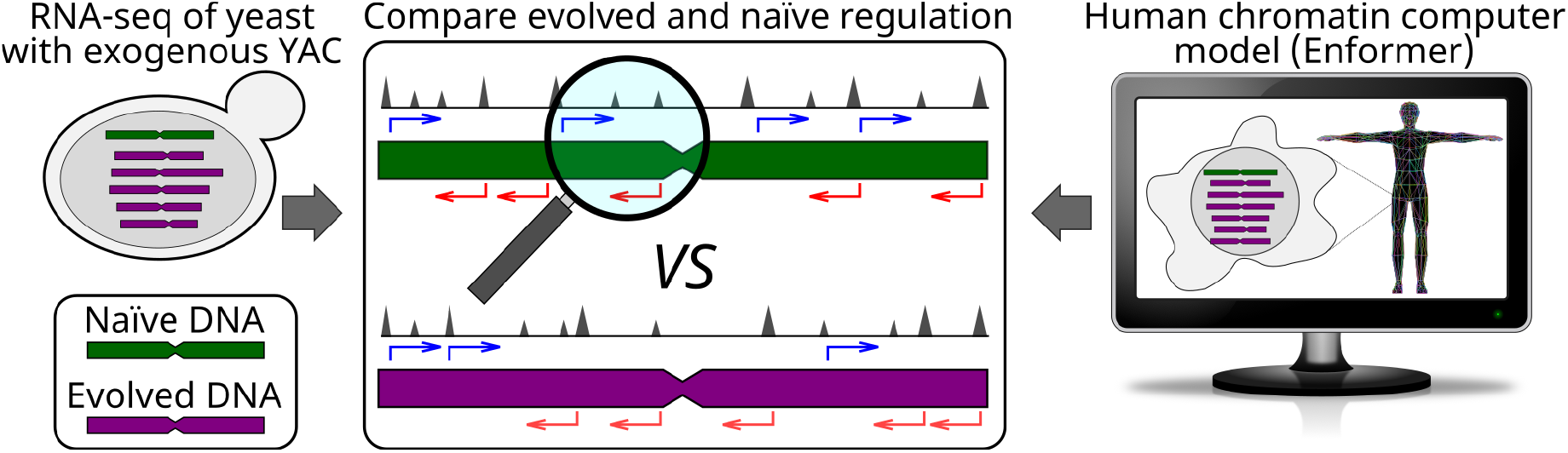
Experimental approach comparing naïve and evolved regulatory activity in human and yeast. For yeast, we performed RNA-seq on yeast harboring exogenous DNA (a human YAC), and compared the transcriptional properties of naïve (exogenous YAC) and evolved (endogenous genome). For humans, we used Enformer (37) to predict chromatin marks on both naïve (randomized sequences) and evolved (human genomic) DNA.

However, naïve transcripts are frequently transcribed on both DNA strands simultaneously, and do not achieve the same extremes of expression as evolved transcripts. Using the Enformer deep learning model, which predicts chromatin features from DNA sequence, we examined the activity of random DNA in human cells and tissues. Similar to yeast, humans are predicted to produce extensive regulatory activity from naïve DNA, but this activity is dependent on preservation of local dinucleotide content. Chromatin marks associated with active regulation are predicted to co-occur similarly in naïve sequences and evolved sequences, but achieve higher extremes in evolved DNA, and regulation of naïve DNA appears to be more cell type specific than evolved DNA. Overall, our results suggest that much of the biochemical activity seen in genomes is expected in the absence of selection.

## Results

### Abundant and diverse transcription of naïve DNA

In order to test the transcriptional activity of evolutionarily naïve DNA, we performed strand-specific poly-A+ RNA-seq of a yeast strain harboring ∼760 kb of exogenous (human) DNA in a yeast artificial chromosome (**YAC**) (33). Since human and yeast are separated by ∼1 billion years of evolution (34,35) and have distinct core promoter structures (36), we expect human DNA to be a good proxy for evolutionarily naïve DNA. Because the exogenous DNA and yeast genomic DNA are being expressed within the same cells, any differences in the transcriptomes they each produce can be attributed to their distinct origins.

Many properties of the naïve and evolved transcriptomes were similar in yeast, indicating that the yeast cell is interpreting the human DNA in a yeast-like way. While the YAC contained the human GNAT3, SEMA3C, and CD36 genes, the transcripts produced by yeast at these loci are distinct from those observed in humans (**Fig. 2A,B; Supplementary Fig. 1**). In order to get an unbiased view of the transcriptome, we called transcripts *de novo* using Cufflinks (38), and compared evolved and naïve transcripts. The density of transcripts was approximately the same (0.42 and 0.47 transcripts/kb for evolved and naïve, respectively), and transcripts were approximately the same length on both evolved and naïve DNA (2011 vs 2108 bp; two sided Wilcoxon rank sum p=0.3; **Fig. 2C**). However, evolved transcripts were expressed more highly on average than naïve transcripts (158 vs 35 FPKM; two sided Wilcoxon rank sum p = 0.0003; **Fig. 2D**). In particular, there were many evolved transcripts with extreme high expression levels that were not present in naïve DNA (**Fig. 2D**), indicating that evolution must optimize to get very high expression levels. Splicing, which is rare in the yeast genome (306/5028 transcripts spliced), was about half as common in naïve DNA (10/356 transcripts spliced; 6% vs 2.8%; two sided binomial p=0.007), with naïve intron sequences closely matching known splicing motifs (39) (**Supplementary Fig. 2**). Overall, naïve DNA produces a strikingly similar transcriptome to that of the evolved yeast genome.

**Figure 2:**
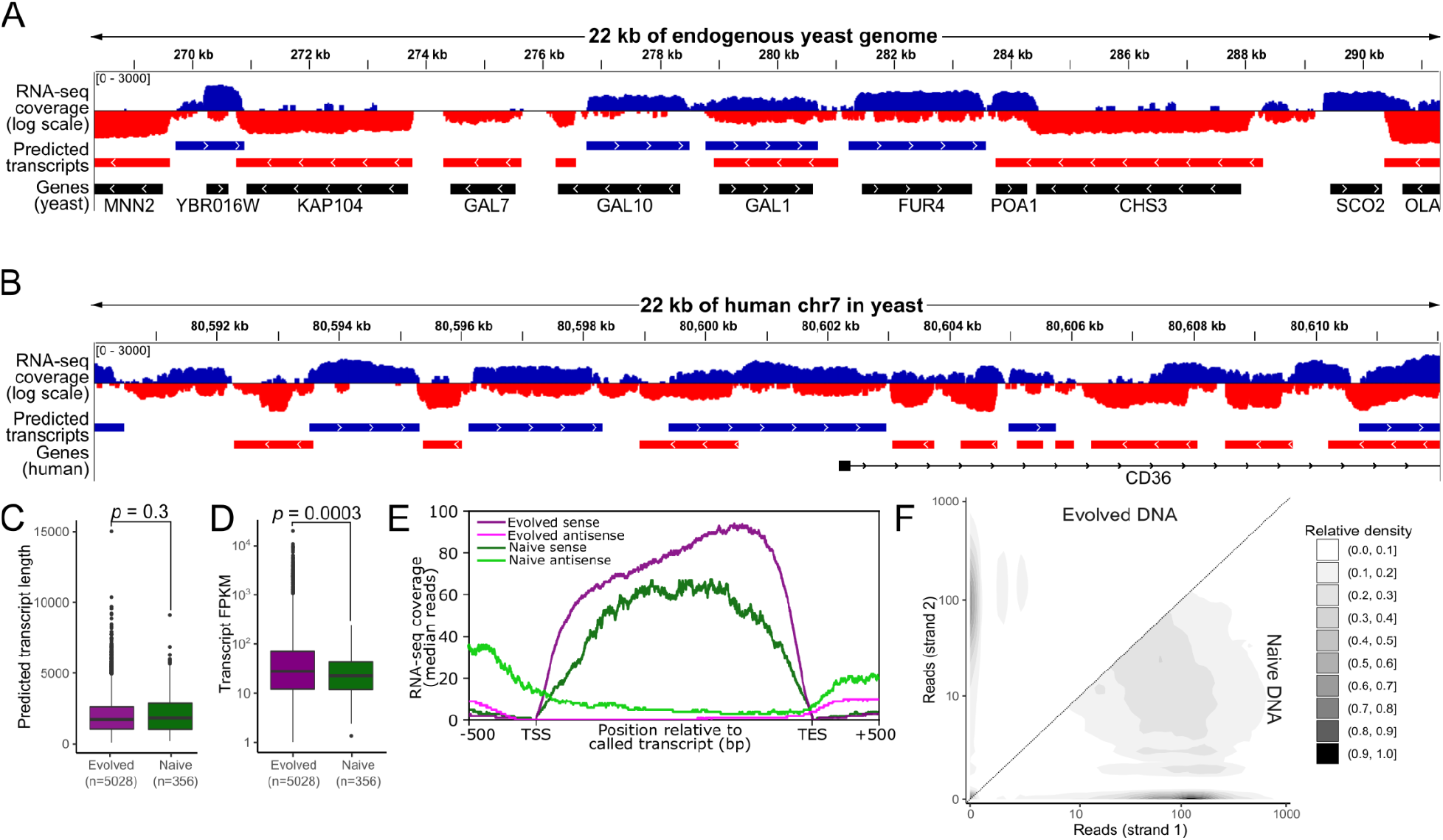
Evolutionarily naïve DNA includes abundant and heterogeneous transcripts in yeast. (**A,B**) Genome browser shots of (**A**) the Gal1-10 locus from the evolved yeast genome and (**B**) part of human chromosome 7, expressed in yeast. Blue=sense transcription; red=antisense transcription (both log scale), with predicted transcripts indicated (red and blue), and known genes indicated in black. (**C**) Boxplot of lengths (y-axis) for evolved and naïve transcripts (*x*-axis and colours). (**D**) Boxplot of expression levels (FPKM; *y*-axis) for evolved and naïve transcripts (*x*-axis and colours). (**E**) Metagene profiles of the expression level (*y*-axis) distribution across the regions containing transcripts (*x*-axis) in evolved and naïve DNA for sense and antisense transcription (colours). (**F**) Contour plot showing expression on one strand relative to the other (*x*- and *y*-axes) for evolved (upper left triangle) and naïve (lower left triangle) DNA.

### Yeast genes have evolved coherent transcript structures

The evolved transcriptome has a much more coherent transcript structure, with more untranscribed bases in between transcripts, and less antisense expression compared to the naïve transcriptome. Specifically, the median RNA-seq coverage across predicted transcripts originating from naïve DNA has substantially more antisense transcription than evolved transcripts across its length (**Fig. 2E**; **Supplemental Fig. 3**). Because of the abundant expression in naïve DNA (**Fig. 2B**), which may impact our ability to predict transcripts, we verified this result by inspecting RNA-seq coverage at base pair resolution. While the evolved DNA was mostly expressed only on one DNA strand, many bases of the naïve DNA were expressed on both DNA strands (e.g. 14% *vs* 40% of bases transcribed at >=10 reads on both strands), and, overall, naïve DNA was expressed on the minor strand at significantly higher levels (Kolmogorov Smirnov P< 2.2 × 10^−16^; **Fig. 2F**; **Supplementary Fig. 4A**). This also revealed that the evolved genome had ∼6x more DNA that was not expressed at all (3.4% *vs* 0.6%) and reached significantly higher expression levels (**Supplementary Fig. 4B**). Thus, the yeast genome evolved to produce coherent transcript structures, minimizing antisense expression and including more of the extremes of expression (both high and low).

### Dinucleotide-dependent biochemical activity in mammals

Creating and delivering synthetic or exogenous DNA to human cells is expensive, laborious, and limited to ∼200 kb (40). Meanwhile, the gene density of the human genome is much lower than yeast (∼0.64 protein coding genes per 100kb), severely limiting our ability to create an empirical regulatory null hypothesis in humans. Instead, we opted to define a regulatory null hypothesis for humans using the deep learning model Enformer (37), which predicts chromatin marks from DNA sequence alone, enabling us to easily and quickly estimate regulatory activity across many sequences. Here, we focused our analysis on several chromatin marks associated with regulatory function, including DNase hypersensitivity (associated with open chromatin at promoters and enhancers), CAGE (associated with transcript initiation by RNA polymerase II), H3K4Me3 (associated with promoters), H3K4Me1 (associated with enhancers), H3K27Ac (associated with both enhancers and promoters), and H3K27Me3 (associated with polycomb repressive complex 2 repression and poised regions) (41).

We found that randomized DNA sequences are predicted to be active in human cells, but require that local dinucleotide content is preserved. Organismal dinucleotide biases are stable across evolutionary time (42) and the human dinucleotide content is highly biased, for instance, with ∼4x as many GpCs as CpGs (43). To test the degree to which base bias influences predicted regulatory activity, we predicted chromatin marks using Enformer on each of 1000 evolved (endogenous genomic) loci (from the Enformer test set) and 1000 randomly generated 200 kb DNA sequences. We tested randomization in seven different ways, including randomly selecting bases from the human genomic average, and shuffling genomic sequences at either the mono-, di-, or trinucleotide level, either globally (across the entire sequence) or locally (over a sliding window of 100 bp (mono- and dinucleotide) or 200 bp (trinucleotide); **Methods**). We confirmed that shuffled sequences had little sequence similarity to the originals (**Supplementary Fig. 5**). First, we looked at predicted DNase I signal in human pluripotent stem cells (**hPSCs**), and found that preservation of local di- or trinucleotide content was needed to produce sequences with regulatory activity approaching evolved sequences (**Fig. 3A,B**).

**Figure 3:**
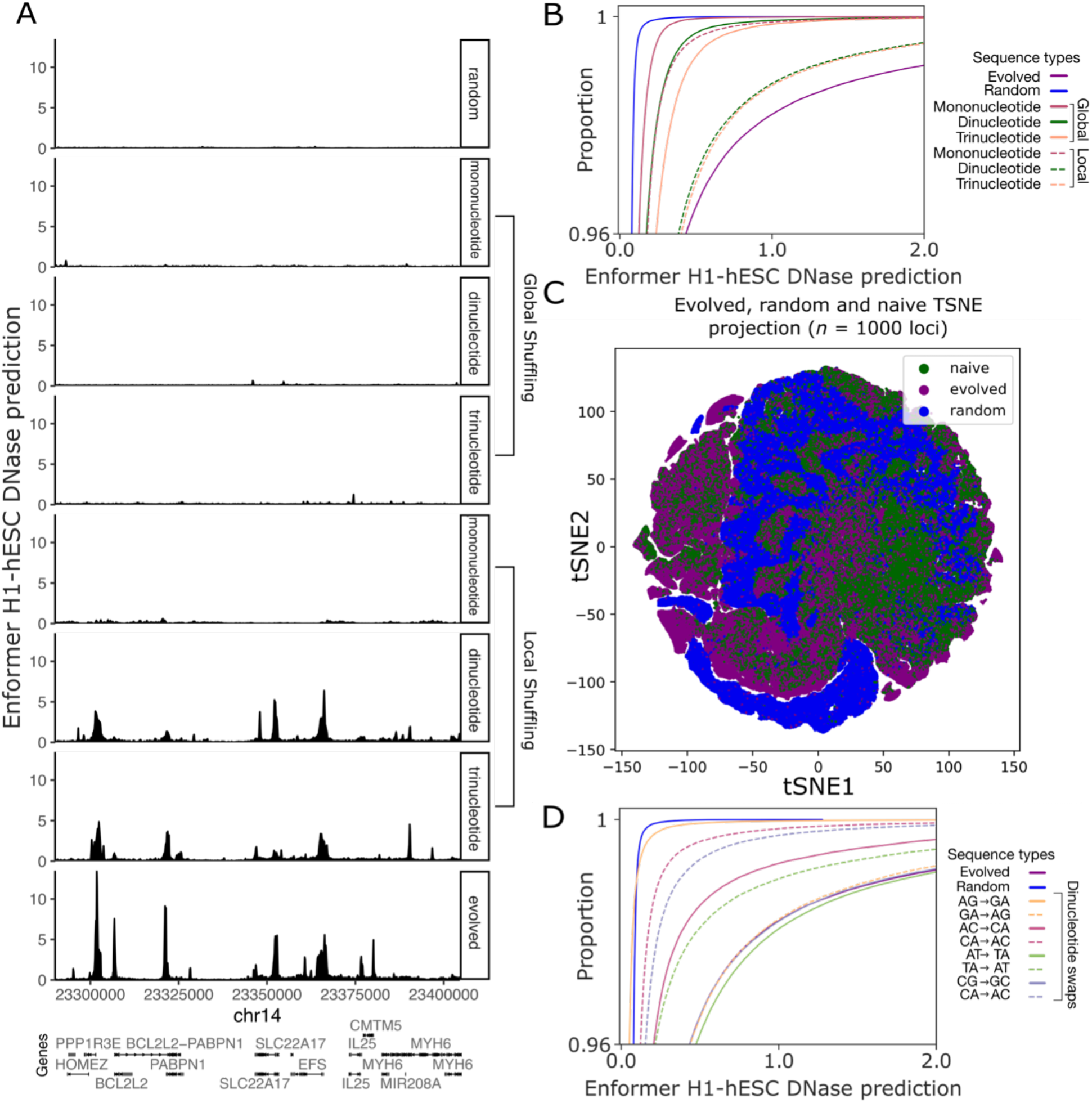
Enformer predicts that locally dinucleotide-matched DNA has regulatory activity. (**A**) Genome browser shot of a locus and randomized counterparts. All tracks correspond to predicted DNase I signal in human Embryonic Stem Cells (hESCs; *y*-axes), with genomic or random sequence location (*x*-axis) and gene annotations corresponding to the genomic sequence. (**B**) The cumulative distribution (*y*-axis; top 4%) of Enformer DNAse I activity in hESCs (*x*-axis) across 1000 115 kb loci for evolved and the different random sequence (colours). (**C**) Clustering of random, local dinucleotide shuffled, and genomic sequences (colours) based on Enformer predictions for H3K27Ac, H3K27Me3, H3K4Me1, H3K4Me3, and DNase I within H1 hESCs. Each point is a 128 bp Enformer prediction bin. (**D**) As in (B), but for random sequences, and evolved sequences with/without different dinucleotide reversals (colours).

Next, we aimed to gauge the relative similarities of random, locally dinucleotide shuffled, and genomic sequences. We predicted regulation-associated chromatin marks (H3K27Ac/Me3, H3K4Me1/3, and DNase I) within hPSCs for purely random, local dinucleotide shuffled, and genomic DNA sequences, and clustered the 128 bp prediction bins based on the levels of these five marks to identify groups of sequences with similar chromatin marks (**Methods**). We found that random sequences largely clustered off by themselves, while genomic and local dinucleotide shuffled sequences co-localized in the vast majority of clusters (**Fig. 3C**), indicating that the local dinucleotide shuffled sequences are predicted to generate similar regulatory activity to genomic sequences, while random sequences are largely distinct.

We hypothesized that the importance of local dinucleotide preservation in producing sequences predicted to be biochemically active could result in part from CpGs, which can be methylated and impact gene regulation, and are highly biased in their abundance and location (43,44). However, we found that this is likely only a partial explanation as reversing CpGs to GpCs (or vice versa) in genomic sequence impacted their abundance as expected (**Supplementary Fig. 6**), but only partly reduced their predicted hPSC DNase I activity (**Fig. 3D**). Meanwhile swapping ApGs and GpAs impacted predicted activity substantially more than CpGs, with activity nearly abolished when ApGs are exchanged for GpAs and the opposite exchange barely affecting the predicted activity (**Fig. 3D**). Because of the strong dinucleotide bias of the human genome and apparent dependance of activity on preservation of local dinucleotide biases, we will use local dinucleotide shuffled genomic sequences as naïve DNA for the remainder of our analyses. This definition of “naïve DNA” can be thought of as aiming to capture the same mutational processes that produced our genomes (via dinucleotide biases) (44), but have not undergone selection for function.

### Naïve human DNA has much of the activity of evolved, but varies by chromatin mark

In order to gauge the relative activity of naïve sequences based on Enformer predicted activities, we compared predicted chromatin activities to *bona fide* peaks called by ENCODE within the same cell types and for the same chromatin marks (14). In order to obtain an estimate for how “real” the Enformer predictions are for each chromatin mark, we identified all bins that overlap ENCODE peaks, and found overall good agreement between predictions and ENCODE peaks (**Supplementary Fig. 11**). To determine the proportion of peak-overlapping bins at each predicted activity level, we determined, for each prediction value threshold, the fraction of threshold-exceeding bins that overlap ENCODE peaks (**Methods**; **Fig. 4A**, cyan curve). In order to obtain the relative abundance of each mark in evolved and naïve sequences for these thresholds, we calculated, for each threshold, the ratio of evolved to naïve sequence bins that exceed the threshold (**Fig. 4A**, yellow curve). We performed this procedure for H1 ESCs across several common histone marks assayed by ENCODE (**Fig. 4A**; **Supplementary Fig. 13)**. When 80% of predicted active bins overlap a *bona fide* peak for that mark, naïve sequences have about half as many open chromatin sites as evolved (**Fig. 4A**), about half as many H3K4me3 sites (promoter-associated), about 1/16th as many H3K4me1 sites (enhancer-associated), and 1/16th as many H3K27ac sites (enhancer and promoter-associated; **Supplementary Fig. 13**). Meanwhile, at this same threshold, H3K27me3, a mark associated with polycomb repression (45), was actually about twice as frequent in naïve DNA (**Supplementary Fig. 13A**). For all marks, the more extreme the predicted value, the rarer naïve DNA achieved these levels, suggesting that selection is required to reach the extreme high end of any chromatin mark (**Fig. 4A, Supplementary Fig. 13**).

**Figure 4:**
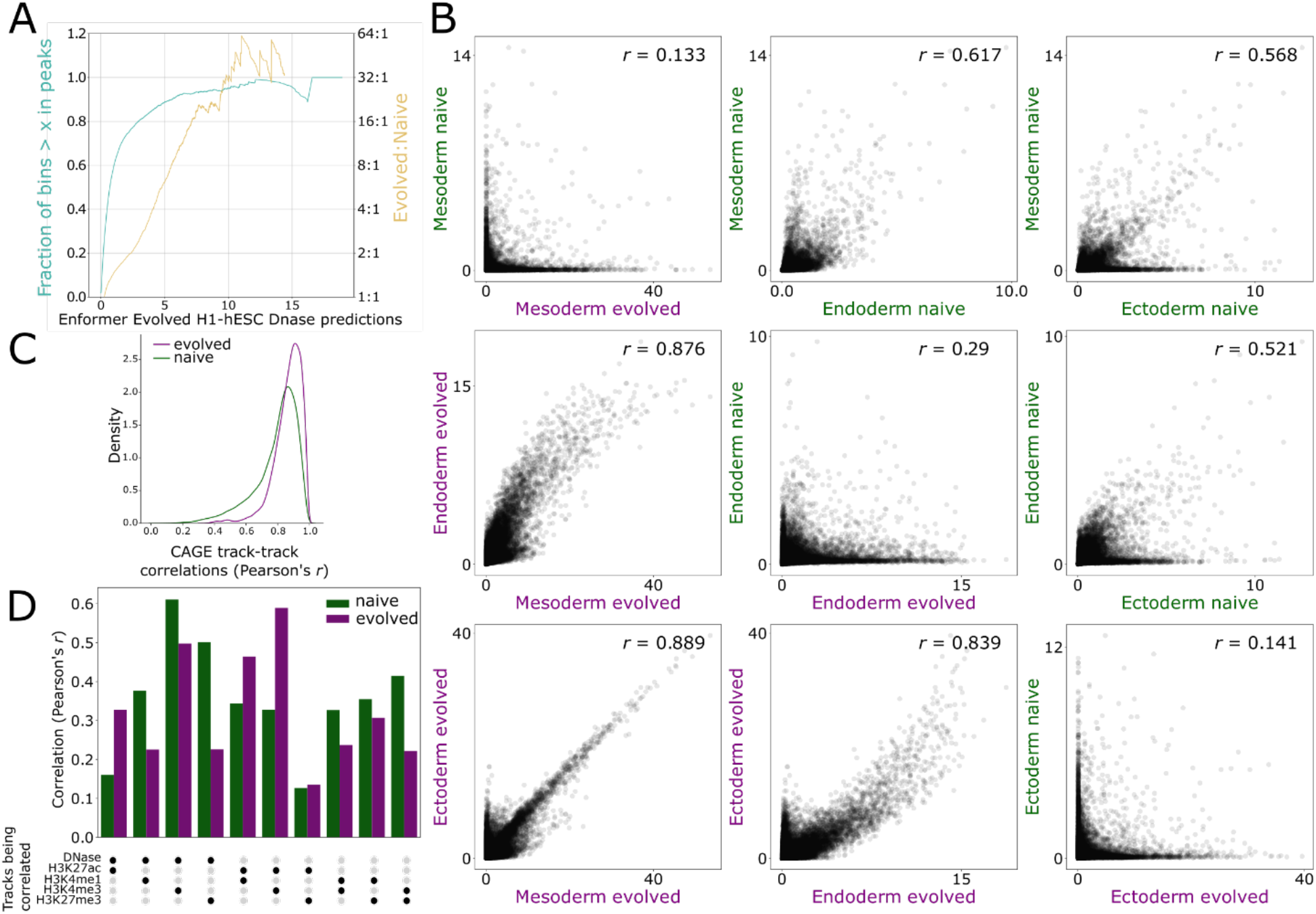
Naïve DNA has about 50% of the activity of evolved sequences, and encodes for cell type specific regulation, with correlated histone marks. (**A**) For each Enformer DNAse I predicted value (*x* axis), the fraction of evolved bins with a score of at least *x* that are within ENCODE DNase I peaks (left *y* axis) and the ratio of evolved to naïve for bins with predicted signals > *x* (right *y* axis). Evolved:Naïve ratio line ends when there are no more naïve bins with a value greater than the *x*-axis. (**B**) Scatter plots comparing predicted DNase I activity in evolved and naïve sequences (*x-* and *y*-axes). Each point corresponds to a 128 bp Enformer prediction bin, with the naïve sequences derived from the evolved sequences (by local dinucleotide shuffling). Ectoderm = foreskin melanocytes, mesoderm = mesenchymal stem cell originated from H1-hESC, and endoderm = hepatocyte originated from H9. (**C**) Density (*y*-axis) of CAGE track-track correlations (*x*-axis) for both naïve and evolved sequences (colours). (**D**) Correlations between prediction tracks for H1 hESCs (*y*-axis) for all pairs of chromatin marks (*x*-axis) for both naïve and evolved sequences (colours).

### Naïve human DNA has cell type specific regulation

Next, we asked whether naïve DNA was predicted to be cell type specific in its activity. First, we predicted DNase I activity in both naïve (local dinucleotide shuffled) and evolved (genomic) sequences for representative differentiated cells from the three major cell lineages that comprise the human body (mesoderm, endoderm, ectoderm). Firstly, we confirmed that naïve sequences (which are dinucleotide shuffled versions of the evolved sequences) do not simply recapitulate the activity of the original sequences (**Fig. 4B**, diagonal, top right to bottom left; Pearson’s *r* = 0.13-0.29), indicating that Enformer has not simply memorized dinucleotide contents of active sequences. Next, we compared predicted DNase I activity between cell types, both within evolved sequences and within naïve sequences, finding that, while DNase I signal was predicted to be quite similar between cell types for evolved sequences (**Fig. 4B**, lower left; Pearson’s *r* 0.84-0.89), they are much more distinct for naïve sequences (**Fig. 4B**, upper right; Pearson’s *r* = 0.52-0.62). Next, we repeated this for all Enformer-predicted CAGE tracks, representing RNA polymerase II initiation. Here, we gauged cell type specificity by the correlation coefficients between pairs of different cell types/tissues, such that lower correlations between tracks result from expression that is more cell type specific (**Methods**). We found that naïve DNA produced more cell-type specific expression than evolved DNA (**Fig. 4C**). This difference was robust even when we selected for only those bins that were predicted to be expressed in at least one cell type (and testing multiple thresholds for “expressed”; **Supplementary Fig. 7**), indicating that this result does not simply result from more noise in predictions at low predicted expression levels in naïve DNA. By both chromatin marks and gene expression, evolved DNA appears to be less cell type specific than naïve DNA in its regulation.

### Naïve human DNA is predicted to have co-occurring chromatin marks

We next asked whether the chromatin marks associated with active regulatory regions in naïve DNA co-occur, which has been suggested to be a marker of selection (2). We predicted chromatin marks within H1 PSCs for both naïve and evolved sequences (**Methods**). First, we co-clustered naïve and evolved sequences based on Enfomer predicted chromatin mark predictions, and found they co-cluster in many regions, including those with co-occurring chromatin marks (**Supplementary Fig. 8**). For a more quantitative analysis, we calculated the correlations between different tracks, with higher correlations resulting from prediction bins that share chromatin marks, indicating co-occurrence. We found that naïve chromatin marks were sometimes more and sometimes less correlated than evolved, but overall, both were quite similar (**Fig. 4D**; **Supplementary Fig. 9**). We repeated this for 11 other healthy cell types for which Enformer could predict these chromatin marks and found largely similar results (**Supplementary Fig. 10**). Overall, naïve DNA has similar correlations between the different histone marks, indicating that this is not a reliable measure of selection.

## Discussion

The surprisingly commonplace biochemical activity of naïve DNA, whether representing DNA from a distantly related organism or randomized DNA, has important implications for our understanding of how genomes evolve. In both yeast and humans, naïve sequences were biochemically active, suggesting that the mere presence of transcription (in yeast) and chromatin marks (in humans) are insufficient to conclude function. Further, naïve DNA in humans is predicted to have cell type specific expression and correlated chromatin marks, indicating that these are not reliable indicators of selection (2), and may instead reflect inherent properties of the transcriptional machinery. In both yeast and humans we found that the extremes of biochemical activity (e.g. histone marks levels, transcription levels) are much more frequent in evolved sequences, indicating that these likely reflect products of selection.

The prevalent basal activity of naïve DNA in yeast and humans may reflect the need for genomes to be evolvable. For instance, an organism whose regulatory machinery is so precise that only functionally important activities occur could be an evolutionary dead end because too many mutations may be required to adapt to a new situation. In contrast, a genome where biochemical activity is relatively easy to achieve is poised for evolutionary optimization since mutations can readily tune an element when it becomes advantageous (22). In most cases this basal activity would be slightly deleterious to the cell due to its associated metabolic costs. Humans are ∼1000x less sensitive to selection than yeast due to differences in our effective population size (46) (∼10,000 for humans (47–49) and ∼10,000,000 for yeast (50)). Consequently humans likely cannot effectively fix genetic variation that downregulates a transcript unless it is very highly expressed or is deleterious beyond its metabolic cost (e.g. the RNA or peptides encoded within interact with other cellular machinery) (7,51). However, mechanisms for global reduction in the cost of this basal activity can evolve. For instance, global noise-reduction pathways, such as the RNA surveillance machinery that can degrade aberrant transcription (7,52), can further reduce the burden of nonfunctional activity, while still providing a pathway for new functional elements to evolve. Further, our finding that naïve DNA tends to be more repressed by default in humans compared with yeast suggests some degree of tuning of this default state to favour genomic silencing in humans. Further, naïve human elements tend to be more cell type specific, limiting their impact to the cells where they are expressed, while leaving a pathway for subsequent functionalization and expanding activity into other cell types. Overall, negative selection will repress detrimental biochemical activities (e.g. via altered regulatory sequence or RNA destabilizing elements (53,54)) until selection is no longer sensitive to them, at which point their evolution will be dominated by random drift (7), also enabling drift in their regulatory function (22). In some cases, these basally active elements can subsequently evolve into functional regulatory elements and genes.

The frequency with which genes evolve *de novo* can be thought of as being captured by a protogene equation (55), describing the probability that a *de novo* gene acquires the necessary features for transcription, translation, and beneficial protein function. We demonstrate that acquiring the signals needed for production of stable transcripts in yeast is simple and is likely not a rate limiting step. Translational signals tend to also be relatively simple and frequently fall along a spectrum of translation efficiency (27). Finally, random proteins can affect fitness with a relatively high frequency (56), although this likely depends on the phenotype being studied. Together, this indicates that *de novo* genes being transcribed, translated, and having a beneficial function may be quite probable in organisms such as yeast. While this de novo gene would initially be far from optimal in terms of transcription, translation, and protein function, its initial expression is a prerequisite to subsequent positive selection. In many cases, *de novo* genes would arise from existing DNA, including from the existing pervasive noncoding transcription in wild type cells (17,57), much of which is antisense transcription (58), which we found to be commonplace in naïve DNA. Further, horizontal gene transfer (59–61) would also be facilitated by transferred genes being expressed to some degree right away even at large evolutionary distances between donor and recipient.

Continued experiments will be necessary to better define the gene regulatory null hypothesis. In yeast, we limited ourselves to a single YAC because the ∼760 kb of DNA and 356 transcripts it yielded made it highly unlikely that our answers would change significantly with more data. Further, we and others have found that random DNA randomized in several different ways produced transcripts in yeast in independent experiments, albeit at smaller scales than that tested here (31,32,62,63), bolstering our yeast findings. Although Enformer’s ability to accurately predict chromatin marks was validated extensively and in multiple ways both by the original authors (37) and subsequently (64), it is still possible that it is to some degree overfit to genomic sequences. Overfitting could lead to underestimation of the regulatory activity of completely random DNA (which is easily distinguished from the endogenous genome even using simple sequence features; **Fig. 3C**). This could similarly lead to underestimation of the activity of local dinucleotide shuffled sequences to some degree. Overestimation of activity of local dinucleotide shuffled sequences by overfitting is unlikely because these sequences are predicted to be cell type specific and in ways that differ from the original genomic sequence that was shuffled. Further, a concomitant study found that reversing but not complementing a human locus (which preserves local mononucleotide content but disrupts local dinucleotide content) destroys the locus’s regulatory activity in mouse embryonic stem cells (62), supporting the importance of local dinucleotide preservation, although their limited scale (∼100 kb) meant that less than one gene was expected in the tested region. A better understanding of the mutational processes that bias genomic sequence would enable better modeling and synthesis of naïve DNA. As technology for synthetic genomics improves, it will be necessary to validate the regulatory null hypothesis with larger scale measurements of naïve DNA in human cells, and to identify the cellular and sequence determinants that predict biochemical activity.

## Acknowledgements

We thank The Centre for Applied Genomics and the Steve Scherer lab for the human chromosome 7 YAC, and BRC-seq for assisting with RNA-seq. This research was supported by the Natural Sciences and Engineering Research Council of Canada (RGPIN-2020-05425), the Stem Cell Network (ECR-C4R1-7), and the Canadian Institute for Health Research (PJT-180537). This research was enabled in part by support provided by WestGrid, Compute Canada (www.computecanada.ca), and Advanced Research Computing at the University of British Columbia.

## Author Contributions

IL and XEC created sequence datasets, set-up the Enformer code and performed Enformer analysis. AMR and ALS worked on setting up Enformer code. CJ performed the experiments and analyzed the yeast data. CGD conceived the project and supervised the research.

## Methods

### Yeast strains and culture conditions

The YAC HSC7E1068 -— originally constructed by the Scherer lab as part of the human chromosome 7 specific YAC library (65) — was obtained from the Centre for Applied Genomics (TCAG) (Toronto, Ontario) as a yeast extract peptone dextrose (YPD) stab. Based on the RNA-seq data, the YAC comprised from 80,372,600 to 81,132,754 of human chromosome 7 (hg38), which was consistent with the previously mapped location. To select for yeast containing the YAC, yeast were streaked onto synthetic defined (SD) -URA plates containing adenine sulfate (Sunrise Science, USA) and were grown at 30 °C for two days. Colony PCR was performed on red colonies to confirm the presence of the YAC with primers E1068F (5’-CGCTGAGGACAACACAGTCT3’) and E1068R (5’-ACACTGTCTGGAAAGCTCGG-3’) using a protocol described previously (66). These primers were designed to amplify a portion of *CD36* - a gene reported to be found in HSC7E1068 (67), which was later confirmed by our RNA-seq results.

### Yeast YAC RNA-seq experiment

Yeast colonies containing HSC7E1068, as confirmed by colony PCR, were grown in liquid SD-URA to an OD_600_ of ∼0.3 - 0.6. Total RNA was extracted from liquid yeast cultures using the hot acid phenol method, described previously (68). Genomic DNA was removed from RNA using Turbo DNase following the manufacturer’s instructions (Invitrogen). Extracted RNA was prepared for strand specific sequencing by the Biomedical Research Centre Sequencing Core (UBC, Vancouver) using the NEBNext Ultra II Directional RNA Library Prep Kit for Illumina and polyA selection (NEB). Libraries were sequenced on an Illumina Miseq machine using 300 cycles, generating 20 million paired end reads.

### Yeast YAC RNA-seq analysis

RNA-seq reads were pre-processed using Trimmomatic v0.36 (69) with the parameters “ILLUMINACLIP:TruSeq3-PE-2.fa:2:30:10:2:true SLIDINGWINDOW:4:10 MINLEN:36”. The ILLUMINACLIP parameter identified and eliminated Illumina adapters specified in the Truseq3 library. Using SLIDINGWINDOW:4:10, reads were scanned in a 4 bp window and trimmed when the average quality dropped below 10. The default parameter MINLEN:36 dropped reads below 36 bases long. After trimming, the quality of the reads were confirmed with FastQC v 0.11.8 (70). A reference genome was constructed by concatenating chromosome 7 from the human reference genome GrCH38 (GRCh38.p13) and the yeast S288C reference R64 (<u>GCF_000146045.2_R64/</u>). RNA-seq reads were aligned to the reference using STAR v2.7.10a (71). First, the reference genome was indexed using the following parameters: --runMode genomeGenerate --sjdbGTFfile --sjdbOverhang 60 --sjdbGTFtagExonParentTranscript Parent -- genomeSAindexNbases 12. Reads were then aligned to the reference using the following parameters: --twopassMode Basic --outSAMtype BAM SortedByCoordinate --outFilterType BySJout --alignIntronMax 5000. PCR duplicates were marked using Picard tools v2.18.7 (Broad Institute).

A genome-guided transcriptome assembly from both evolved and naïve DNA was constructed using Cufflinks v0.17.0 (38). The sorted BAM from the STAR alignment was used as input with the following parameters: --no-update-check --overlap-radius 1 --max-intron-length 5000 -- library-type fr-firststrand. The output GTF file containing predicted transcripts was imported into the integrative genomics viewer (IGV) genome browser (Broad Institute) for visual analysis (**Fig. 2A,B, Supplementary Fig. 1**).

Metagene profiles (**Fig 2E, Supplementary Fig. 3**) of the expression level distribution in evolved and naïve DNA (for sense and antisense transcription) were generated using the deeptools v3.5.1 (72) calcMatrix (“-a 500 -b 500 –binSize 1 –regionBodyLength 2050”, and excluding the rDNA locus (chrXII:451500-469100), plotProfile (“--yMin 0 –yMax 100 – averageType median”), and plotHeatmap ‘--sortRegions descend --sortUsing mean -min 0 -- heatmapHeight 4 --heatmapWidth 8 --whatToShow “heatmap and colorbar” --colorList black,r,darkorange,orange,yellow,lightyellow,white’ with “ -max 2250” for sense expression and “-max 300” for antisense). The predicted transcript annotation .gtf file from cufflinks was used as input.

The density contour plot (**Fig. 2F**) showing expression on one strand relative to the other for evolved and naïve DNA was generated using RNA-seq coverage data from BIGWIG files. The deepTools v3.1.3 bamCoverage module was used to generate BIGWIG files from sense (using parameters: -of bigwig --filterRNAstrand forward --binSize 1) and antisense (using parameters: - of bigwig --filterRNAstrand reverse --binSize 1) strands separately. PyBigWig v 0.3.18 was used to extract coverage information for each position along the genome from BIGWIG files (one file for sense, another for antisense). rDNA from chromosome 12 of the endogenous yeast genome was excluded from this analysis due to its propensity to copy number variation, frequently leading to coverage artifacts.

### Enformer overview

Enformer (37) was used to predict chromatin profiles for human genomic (evolved) and random (naïve) DNA. Genomic sequences were retrieved and Enformer run using tensorflow (v2.4.1), and kipoiseq (v0.5.2). To minimize leakage from the genomic regions used to train Enformer, 1000 regions from the test data set were randomly selected (hg38 coordinates), excluding any sequences that had N’s in the sequence. The length of sequence required for Enformer is 393,216 bp but only 196,608 bp of that are used in predicting the 114,688 bp prediction window. This was confirmed by verifying that the predictions from a 393 kb input sequence do not change if the flanking 98,304 bp on either side is replaced with N’s.

The output from Enformer contains 5313 tracks, each of which is a prediction of a specific chromatin feature for a specific cell type. Each predicted track at each locus is comprised of 896 bins, with each bin covering 128 bp (114,688 bp total). We focused our analysis on DNase-seq, ChIP seq of chromatin activity markers (H3K27ac, H3K4me1, H3K4me3, and H3K27me3) since these are commonly measured and associated with gene regulation. In order to test tissue specificity, we selected 12 cell types (H1 cells and those in **Supplementary Fig. 10**; **Supplementary Table 1**) for which all aforementioned markers are predictable by Enformer. These cell types were selected based on criteria that 1) the cells were not derived from cancer cell lines, 2) the cells were not documented to be treated by any medication, 3) the cells were comprised of cells derived mostly from one germ layer (to avoid predictions that correspond to a mixture of heterogeneous cell types), 4) the cells were young so as to avoid epigenomic changes that might take place over one’s lifetime.

### Generation of random, shuffled, and swapped sequences

Random and randomized sequences were generated in several ways. For completely random DNA, we generated 1000 sequences where bases were drawn randomly from the four possible bases, where G and C were drawn with a 20.5% probability, reflecting the 41% GC content of the human genome (73). The random sequence generation was generated by using python (v3.7.11) and numpy.random (v1.19.5). Shuffled sequences were created using BiasAway (v.3.3.0) (74) and used the 1000 genomic regions being analyzed as input. Global shuffling was done by altering the ‘–kmer’ flag in BiasAway. The local mono and dinucleotide shuffling were done with a window size of 100 bp and a step size of 50 bp, and the trinucleotide shuffling used a window size of 200 bp and a step size of 100bp. We confirmed that sequences had diverged from the originals by global alignment between original and shuffled sequences using EMBOSS stretcher (default parameters; **Supplementary Fig. 5**). Dinucleotides were swapped in eight separate tests (e.g. AC to CA and GT to TG is one test because GT is the reverse complement of AC; AA cannot be swapped because it is the same reversed). For each test, dinucleotides were swapped for each of the 1000 genomic regions of study. For each region, every original instance of a dinucleotide was replaced by its reversed sequence using the command line tool sed in a reverse complement aware way. Note that the original dinucleotide will still be found because it can be re-formed during the swap (e.g. replacing the AC with CA in ACC results in CAC and thus still includes an AC), leading to only a partial loss of the original dinucleotides (**Supplementary Fig. 6**).

### Enformer prediction analysis

Enformer’s DNase I predictions for genomic sequences were compared to random sequences and shuffled sequences, as well as dinucleotide swapped sequences using empirical cumulative density distributions (**Fig. 3B,D**), generated using seaborn’s ecdfplot function (v0.11.2).

To visualize the predictions in 2-dimension we used t-SNE dimensionality reduction from scikit-learn (v1.0.2). The predictions from genomic, random, and dinucleotide local shuffled sequences were normalized using the StandardScaler function from sklearn to correct for scaling differences between the tracks (normalizing each track from the three sequence types together). The resulting matrix had 5 features (the 5 tracks) and 2,688,000 (1000 × 896 × 3) data points (the 128 bp prediction bins across all genomic/random/shuffled loci), and was used as input to t-SNE (**Fig. 3C**). This procedure was repeated, but excluding random sequences to embed genomic and dinucleotide local shuffled sequences together (**Supplementary Fig. 8**) to show localization of each chromatin mark prediction.

To calculate a continuous estimate for how reliable Enformer predictions were, we compared activity predictions for bins with ENCODE peaks. The peak data for each H1-hESC track was downloaded from the ENCODE project (https://www.encodeproject.org) (14). For DNase, the two narrow peak bed replicate files were merged [ENCFF726ZCF, ENCFF810HSN]. For all other histone marks the “replicated peaks file” was used; CHIP:H3K27ac [ENCFF520LUH], CHIP:H3K4me1 [ENCFF711NNX], CHIP:H3K4me3 [ENCFF760ZNE], CHIP:H3K27me3:H1-hESC [ENCFF760MIC]. To merge the ENCODE peaks with the Enformer bins we used *bedtools intersect* (75) to find bins that had at least 10 bp overlap with at least one peak. Enformer predictions clearly distinguished bins in peaks vs those not in peaks (**Supplementary Fig. 11)**. Next, the ECDF function from the statsmodels in python was used to calculate the survival curves (1-ECDF) for both genomic and dinucleotide shuffled predictions for all 5 H1-hESC tracks (**Supplementary Fig. 12)**. For each prediction value, the ratio of these survival curves (evolved vs. naïve) was calculated to gauge the relative abundance of each mark in naïve (local dinucleotide shuffled sequences) and evolved (log2((1-ECDF_evolved)/ (1-ECDF_naïve); **Supplementary Fig. 13** and **Fig. 4A** right *y*-axes). For all tracks except H3K27me3, we ceased drawing the abundance ratio curves when there were no more Enformer predictions for the dinucleotide local predictions (at which point the *y*-axis value is undefined). H3K27me3 had similar maximums (max(naïve) = 83.677, max(evolved) = 83.03234100341797) for both genomic and dinucleotide shuffled.

To gauge cell type specificity using CAGE data, CAGE predictions were Z-score normalized, using the mean and standard deviation calculated from the evolved sequence data to scale both evolved and naïve sequences. The correlations between CAGE tracks were calculated for all possible pairings of CAGE tracks, separately for the evolved (genomic) and naïve (dinucleotide local shuffled sequences) (**Fig. 4B**). To ensure that the lower correlations found in naïve CAGE predictions was not dominated by noise at the low end of predictions, prediction bins were filtered to include only bins where at least one CAGE track surpassed a given Z-score threshold, and testing various thresholds (**Supplementary Fig. 7**).

## Data Availability

All data generated in this study are available at NCBI’s GEO database under accession GSE217781.

## Supplementary Figures

**Supplementary Figure 1:**
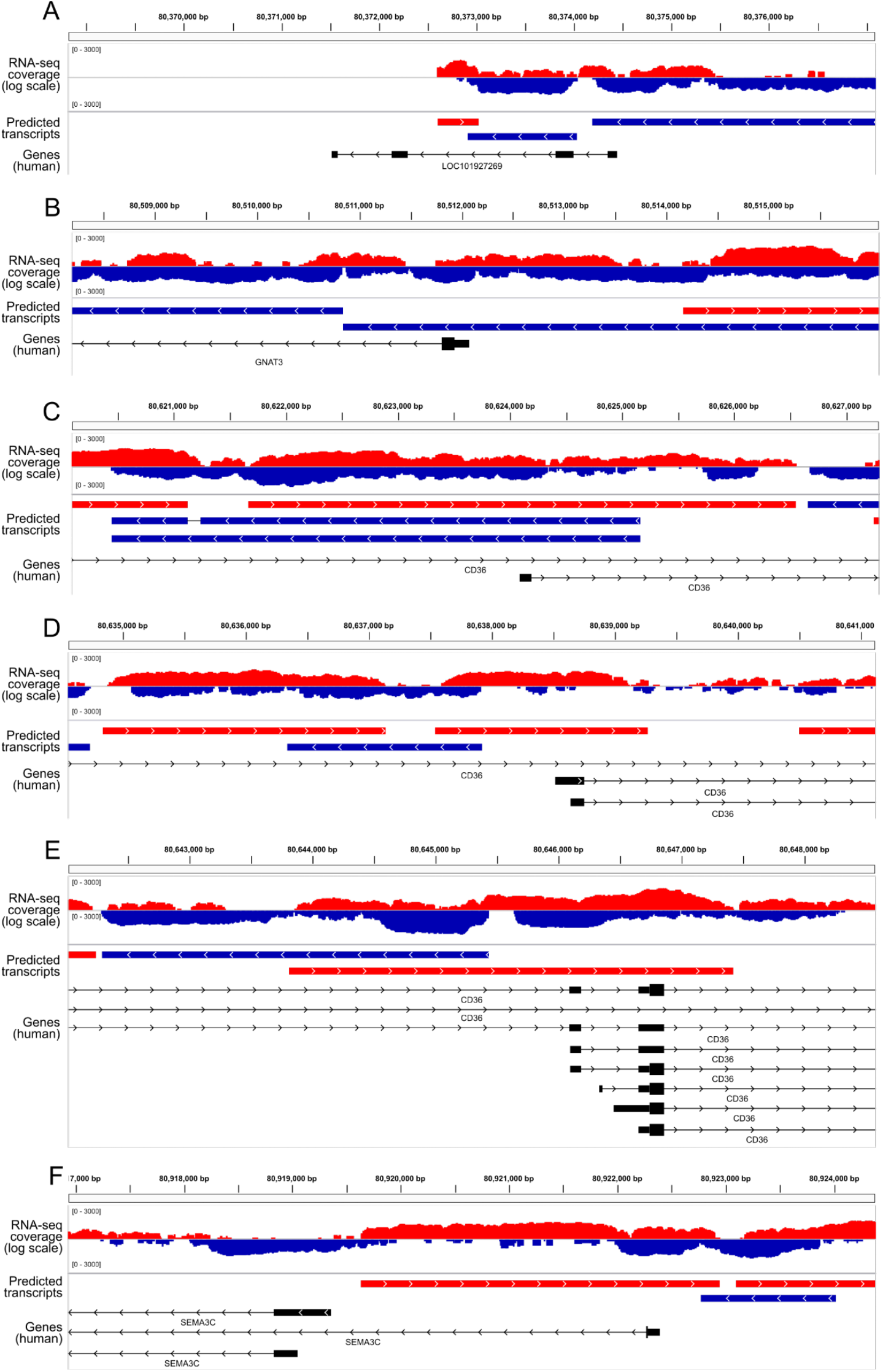
Little correspondence between human promoters and promoter activity in yeast. Yeast RNA-seq coverage for sense (red) and antisense (blue) expression (top), with predicted transcripts (red/blue, as before; middle), and human gene isoforms (refGene; black) are shown below with redundant isoforms collapsed. All refGene TSSs in the tested YAC are shown, including LOC101927269 (which is partly on the YAC; A), GNAT3 (B), CD36 (C,D,E), and SEMA3C (F).

**Supplementary Figure 2:**
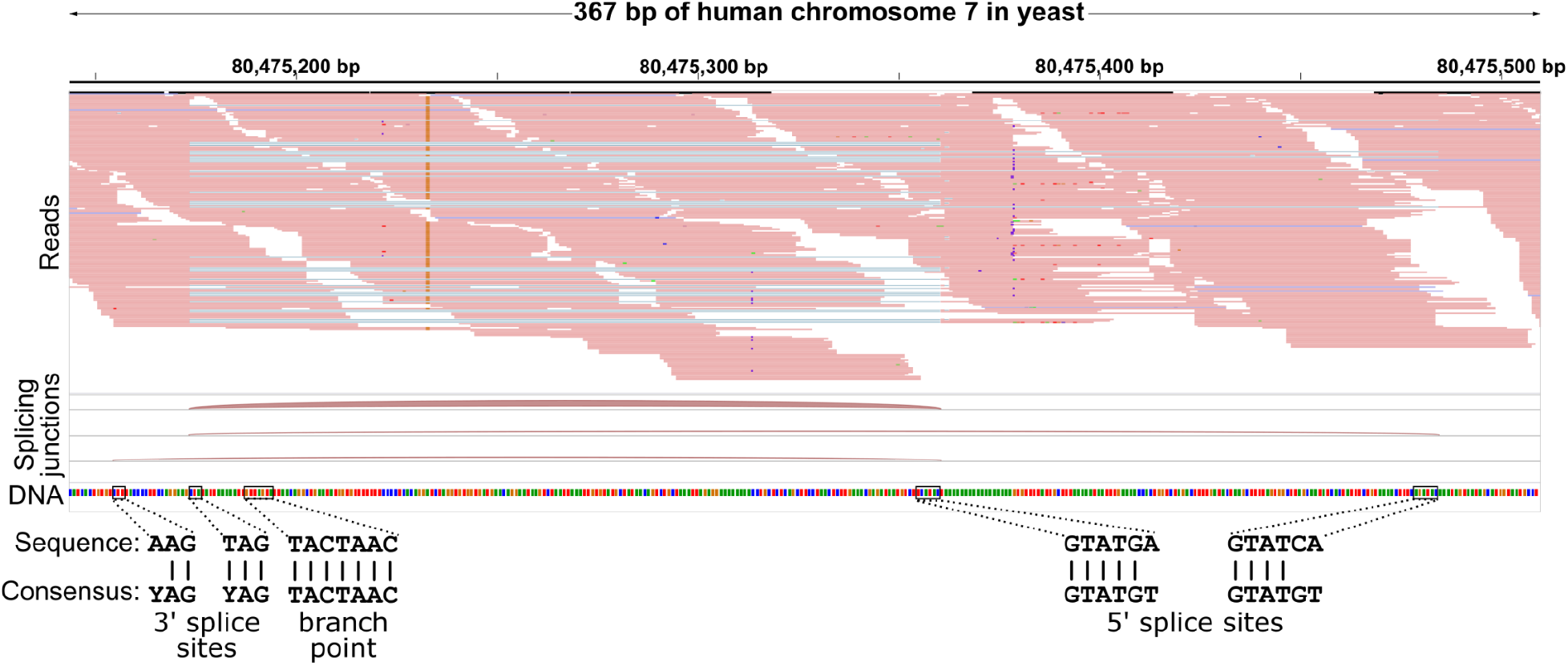
Example spliced naïve transcript. Reads (top) and splice junctions for three isoforms (middle) are shown, with the DNA sequence (colours) at the mid bottom. At the very bottom matches to the known splicing motifs are indicated. The transcript is expressed in the antisense direction.

**Supplementary Figure 3:**
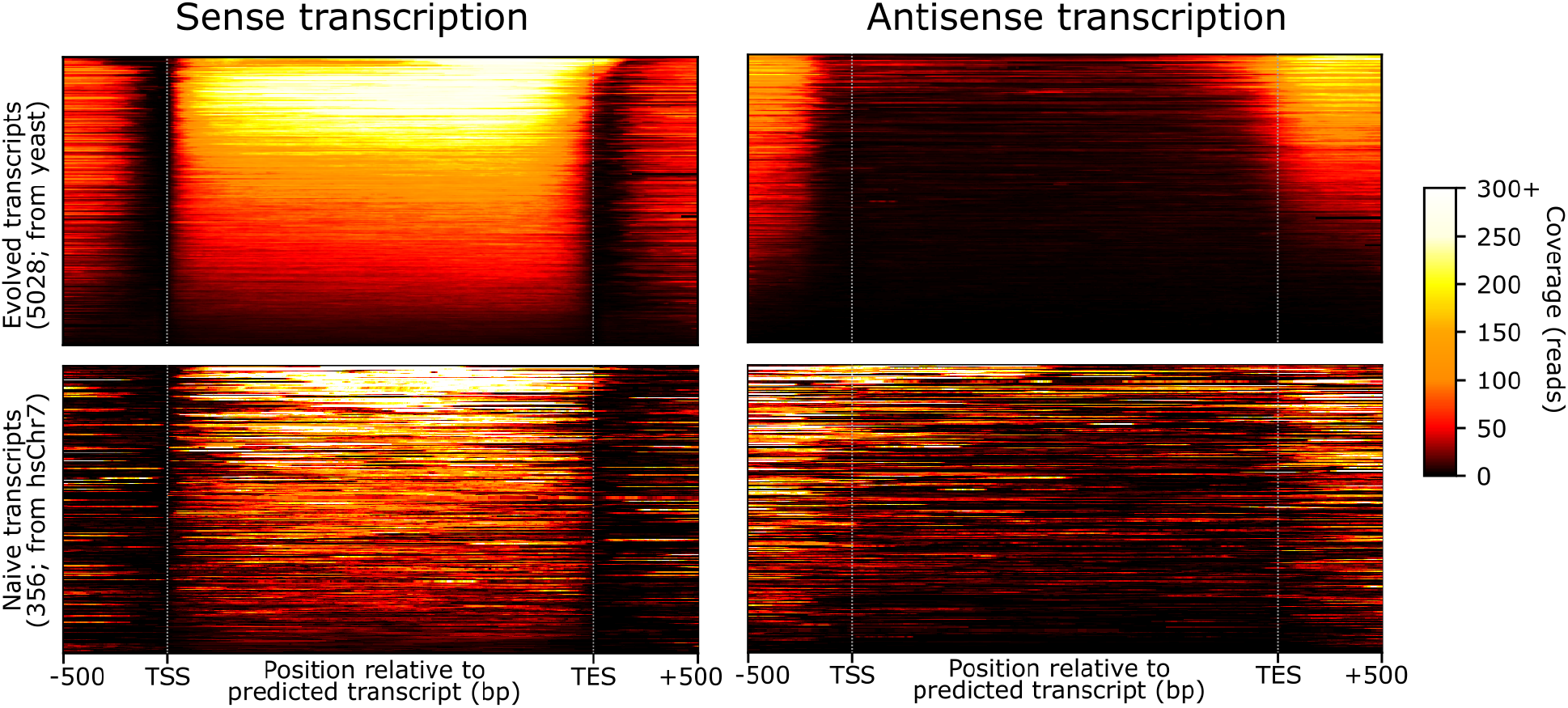
Expression across predicted transcripts for evolved and naïve DNA. RNA-seq coverage (colour) across predicted transcripts (*x*-axes) for predicted transcripts (*y*-axes), for sense (left panels) and antisense (right panels) transcription, and for evolved (upper panels) and naïve (lower panels) transcripts.

**Supplementary Figure 4:**
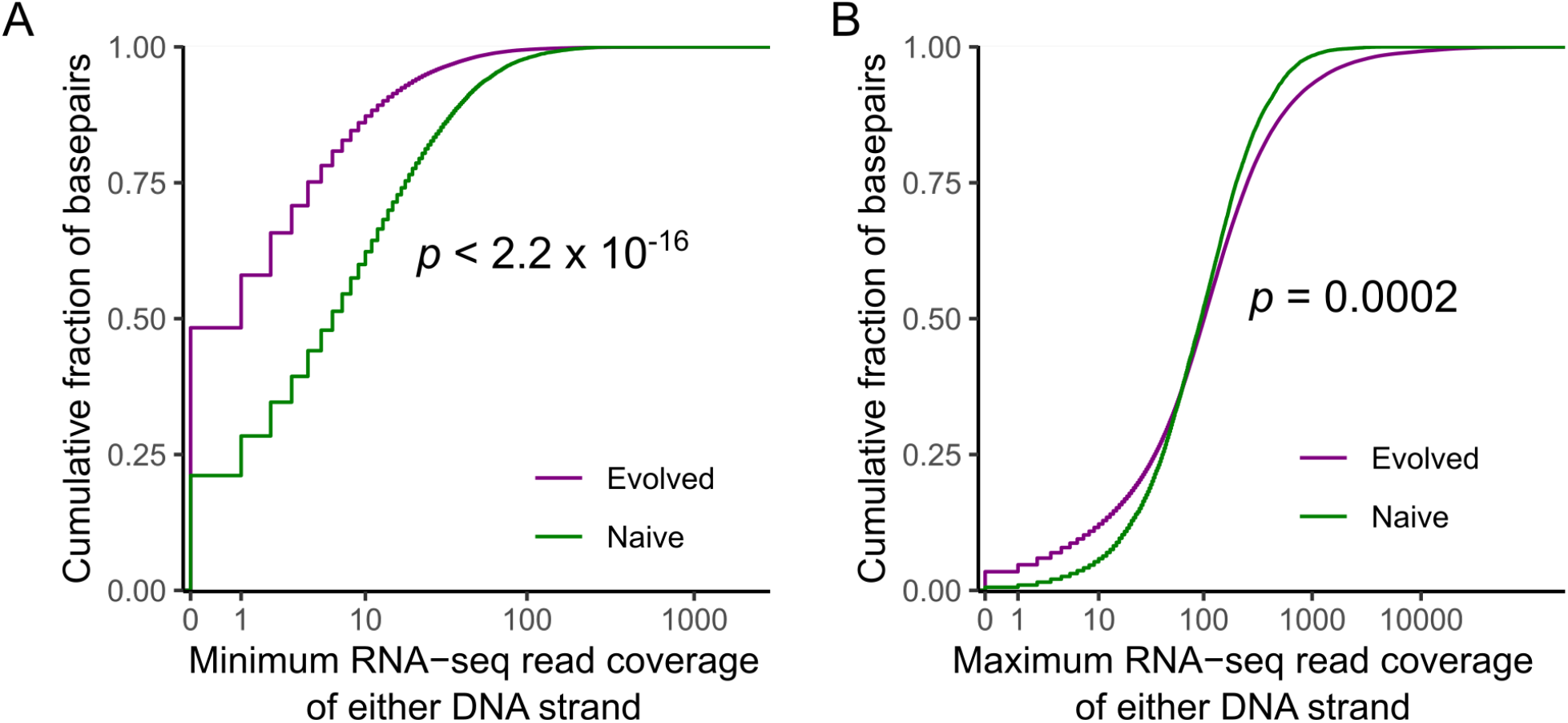
Evolved DNA has less antisense expression, more high expression, and more no expression. (A,B) Cumulative fractions of base pairs (*y*-axes) for each RNA-seq read coverage (*x*-axes), for both naïve and evolved DNA (colours), for (A) the minimum coverage of the two DNA strands or (B) the maximum coverage of the two strands. P-values from Kolmogorov-Smirnov test, using only every 1000^th^ base (to ensure independence between samples); n_Evolved_=12054, n_naïve_=760. Plots include all bases.

**Supplementary Figure 5:**
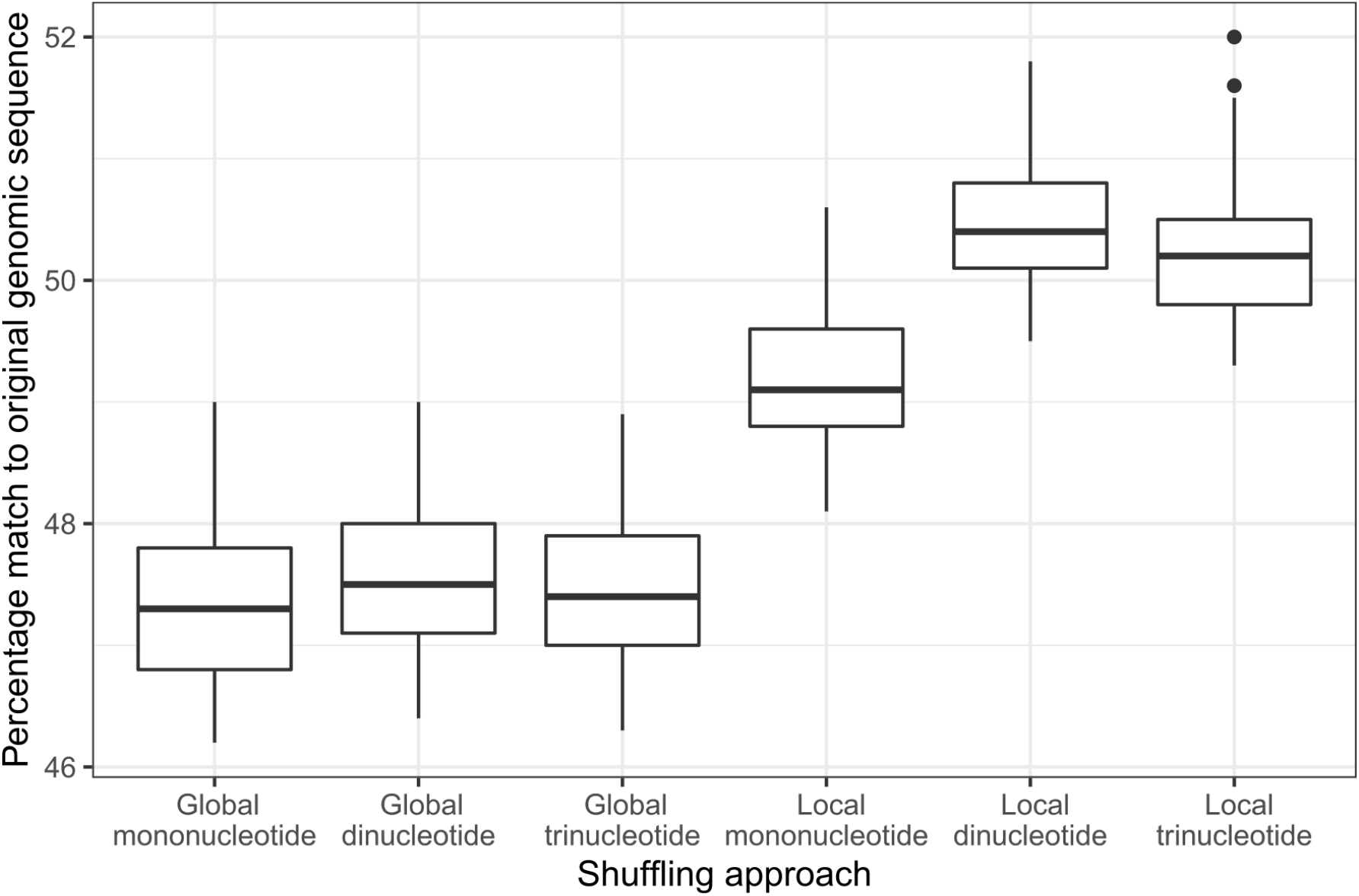
Shuffled sequences bear little resemblance to originals. Percentages of matching bases for each shuffled sequence relative to the original genomic sequence (*y*-axis) for each shuffling approach (*x*-axis). Outliers likely represent sequences containing abundant simple repeat elements.

**Supplementary Figure 6:**
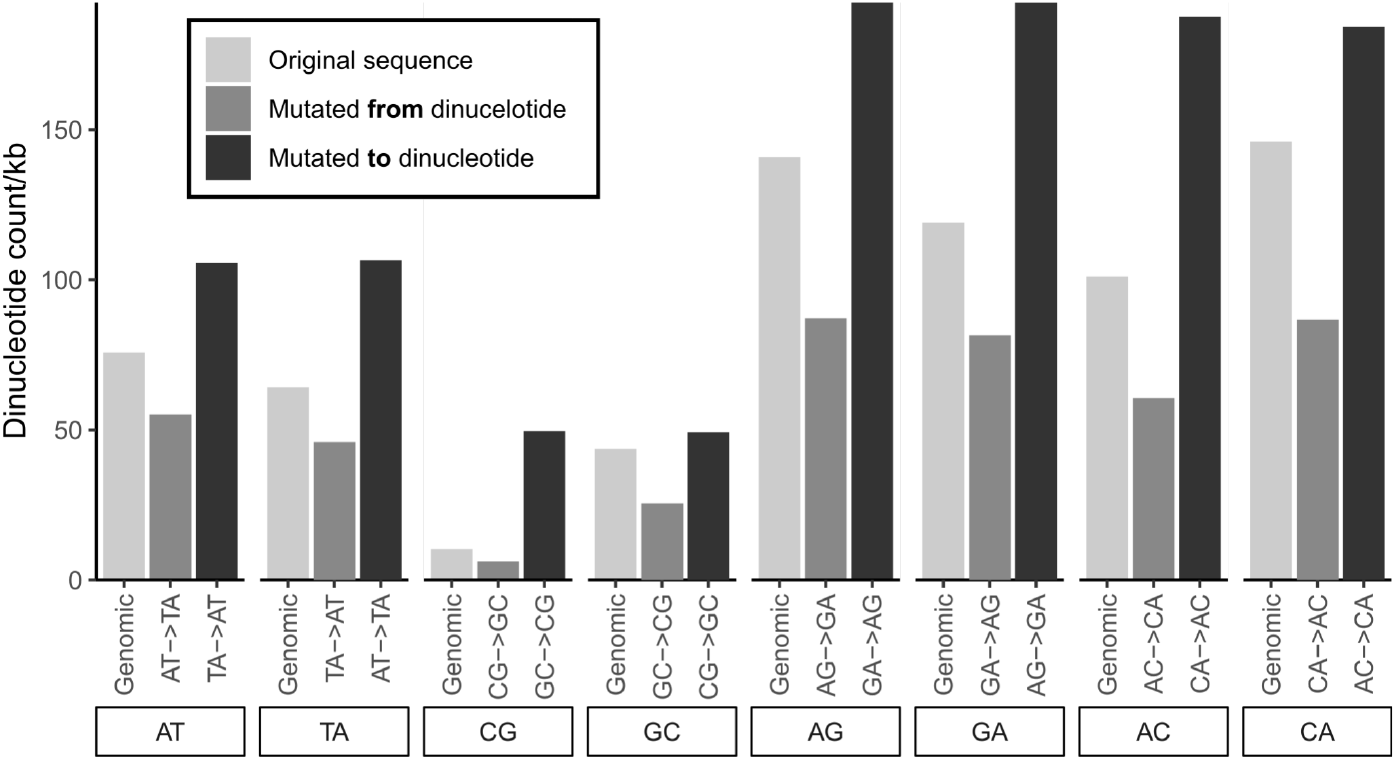
Dinucleotide frequencies before and after dinucleotide mutations. Dinucleotide counts per kb (*y*-axis) for each dinucleotide (*x*-axis, lower labels) for original genomic sequences and each dinucleotide mutation (*x*-axis, upper labels), with colours indicating the nature of the mutations.

**Supplementary Figure 7:**
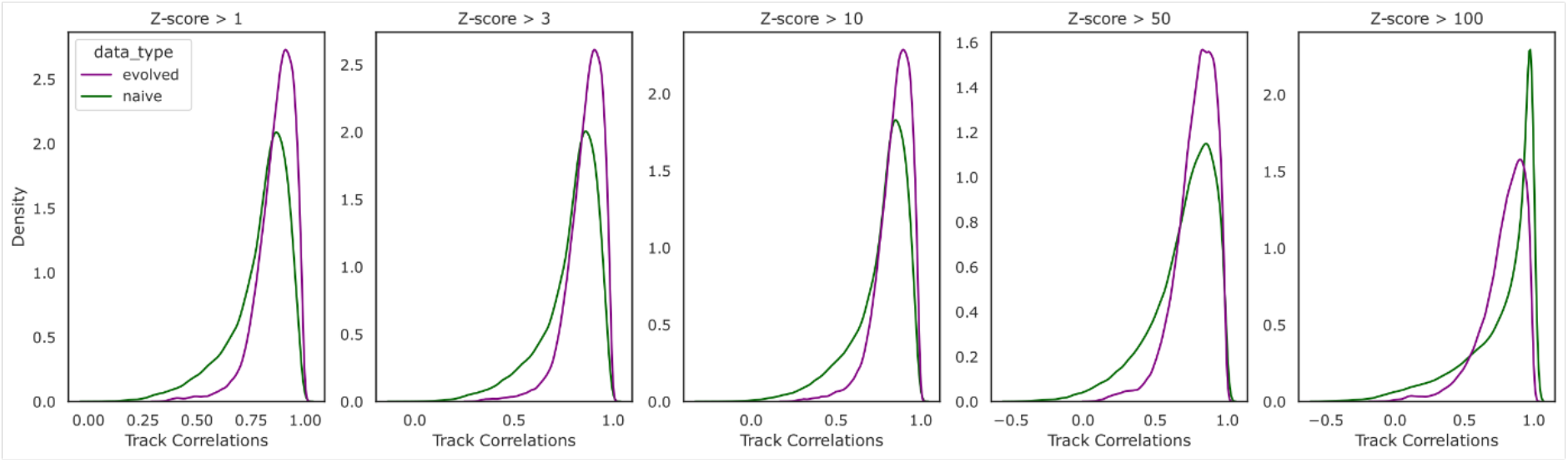
Density plots of genomic and dinucleotide local shuffled CAGE track correlations. Filtering values at different z-score thresholds (keeping highly expressed bins) shows the same distributions up till Z > 50.

**Supplementary Figure 8:**
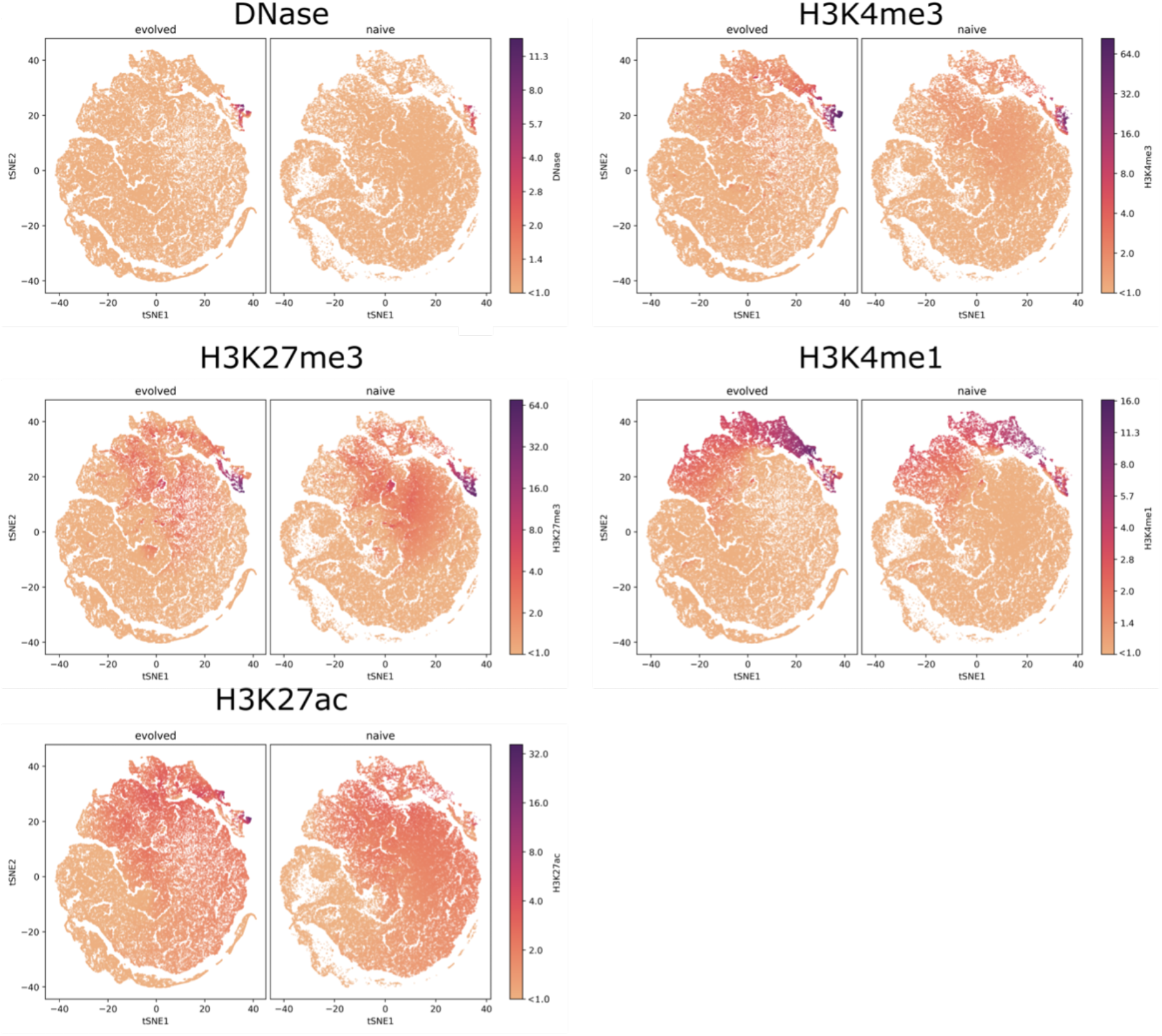
TSNE plots split into points from evolved in first panel and naïve in the second panel. Each point is coloured by the track value predicted from Enformer. We find shared localization of highly predicted marks in evolved and naïve clustered regions.

**Supplementary Figure 9:**
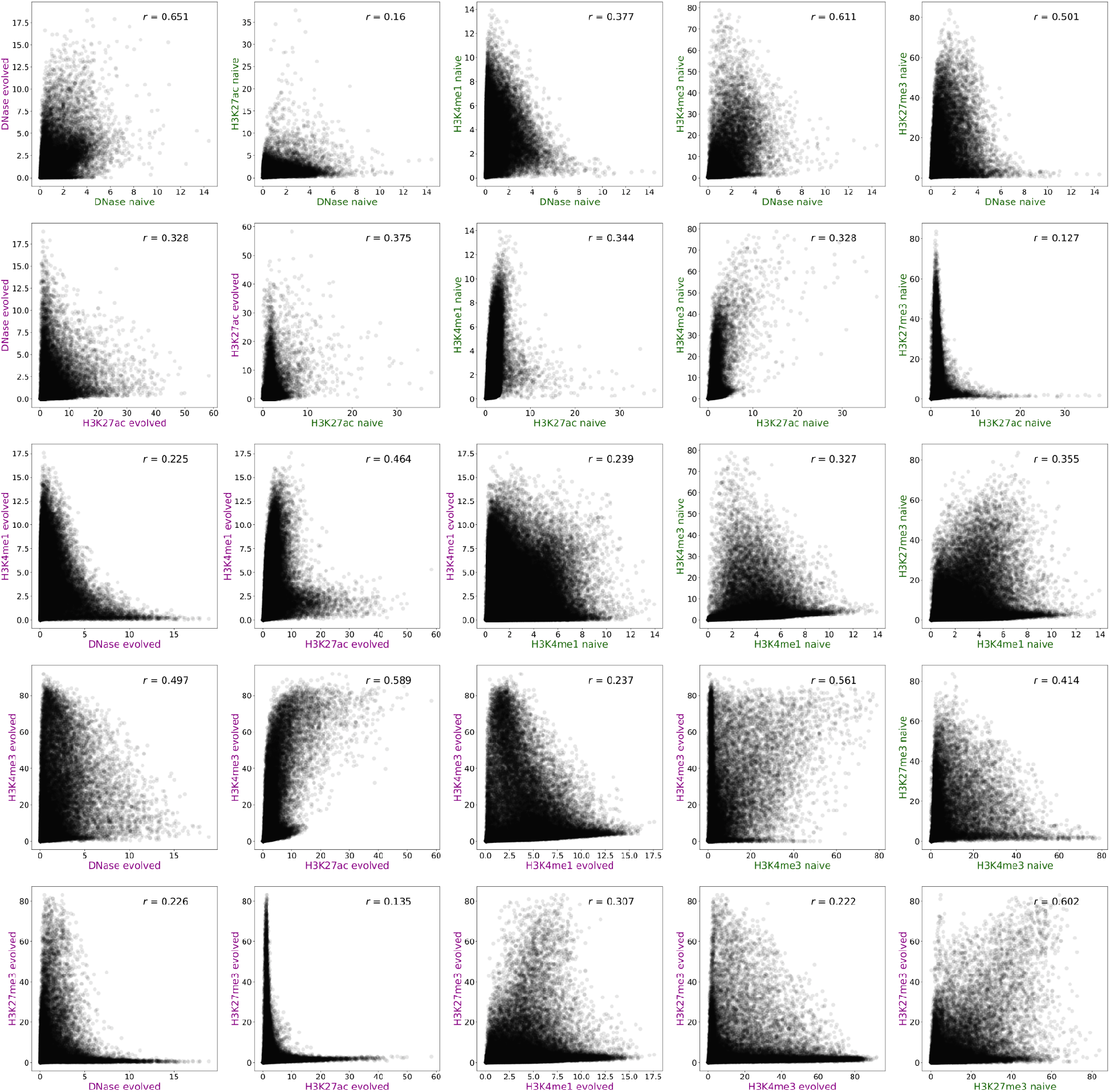
Scatter plots of predicted values for all 1000 regions between all pairs of the 5 tracks. Each scatter plot shows the co-occurrence of the predicted values between two tracks. Scatter plots on the diagonal show the comparison between the naïve and evolved predicted values for the same track.

**Supplementary Figure 10:**
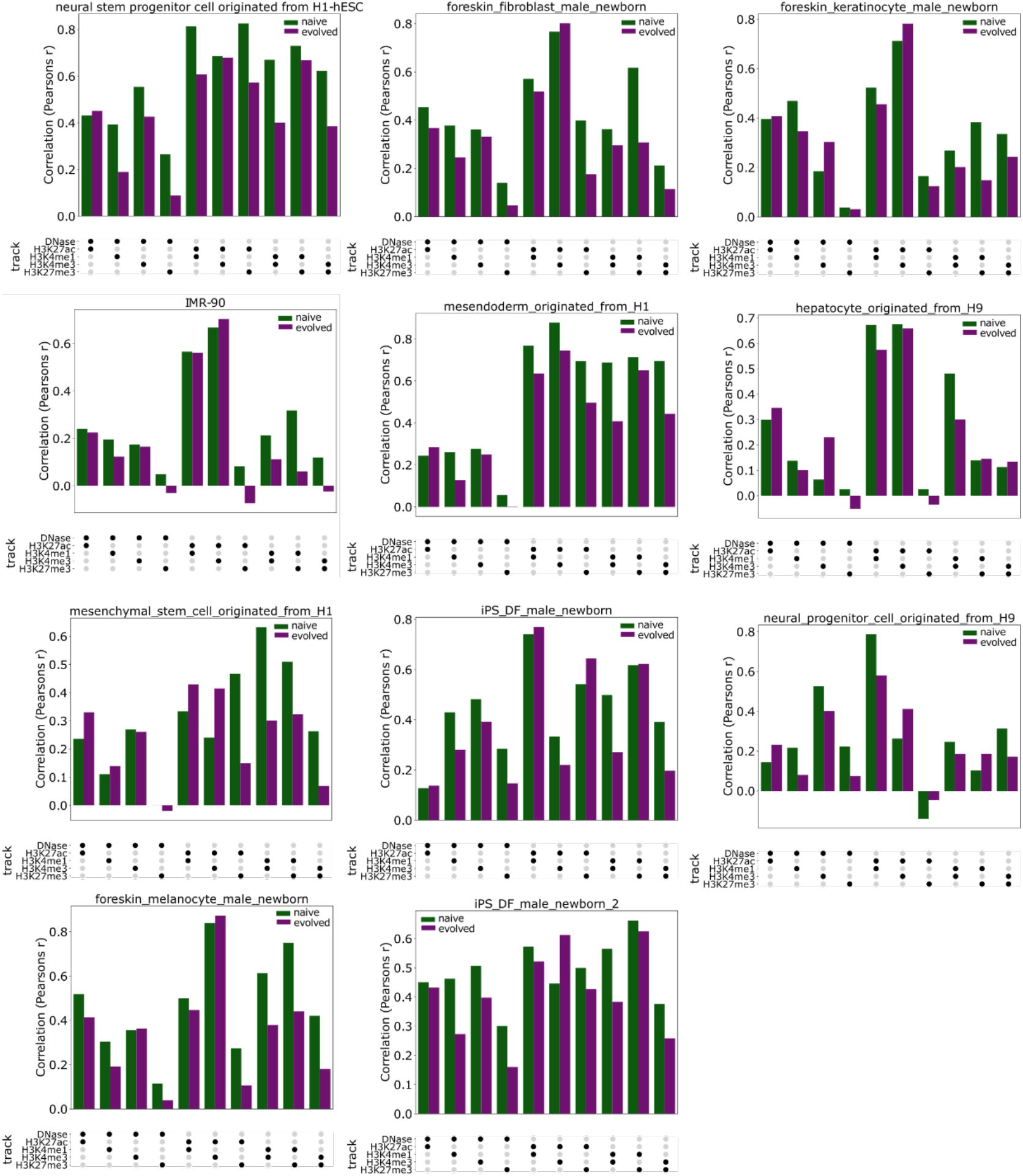
Similar chromatin mark correlations across cell types. Pearson correlation coefficients (*y*-axes) for different pairs of chromatin marks (*x*-axes) for both naïve and genomic sequences (colours) across 11 human cell types (indicated above each graph).

**Supplementary Figure 11:**
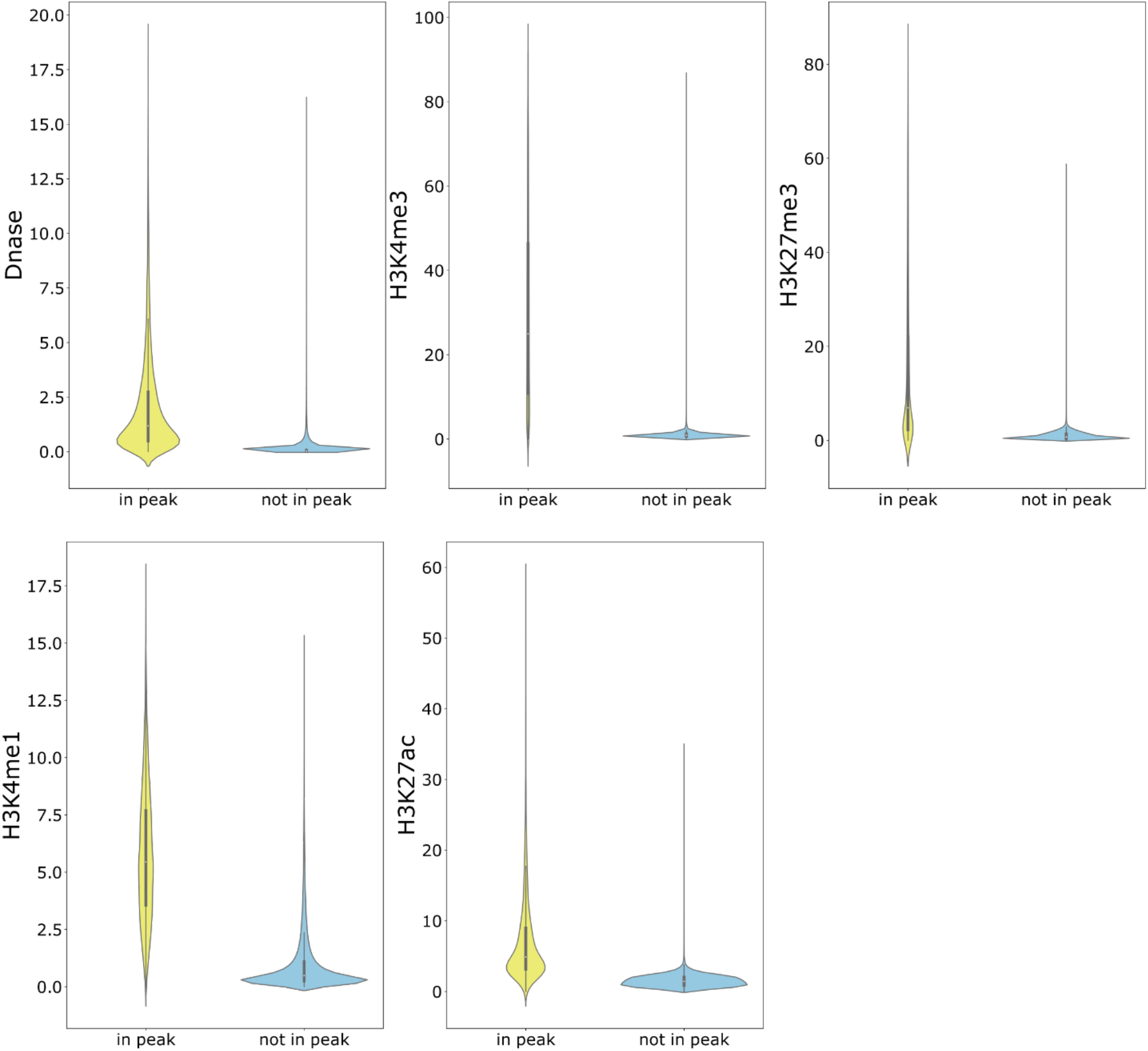
Bins in peaks have higher predicted values compared to those not in peaks. Distribution of Enformer predicted values (*y*-axes) for bins that overlap an ENCODE peak by at least 10 bp or do not (*x*-axes).

**Supplementary Figure 12:**
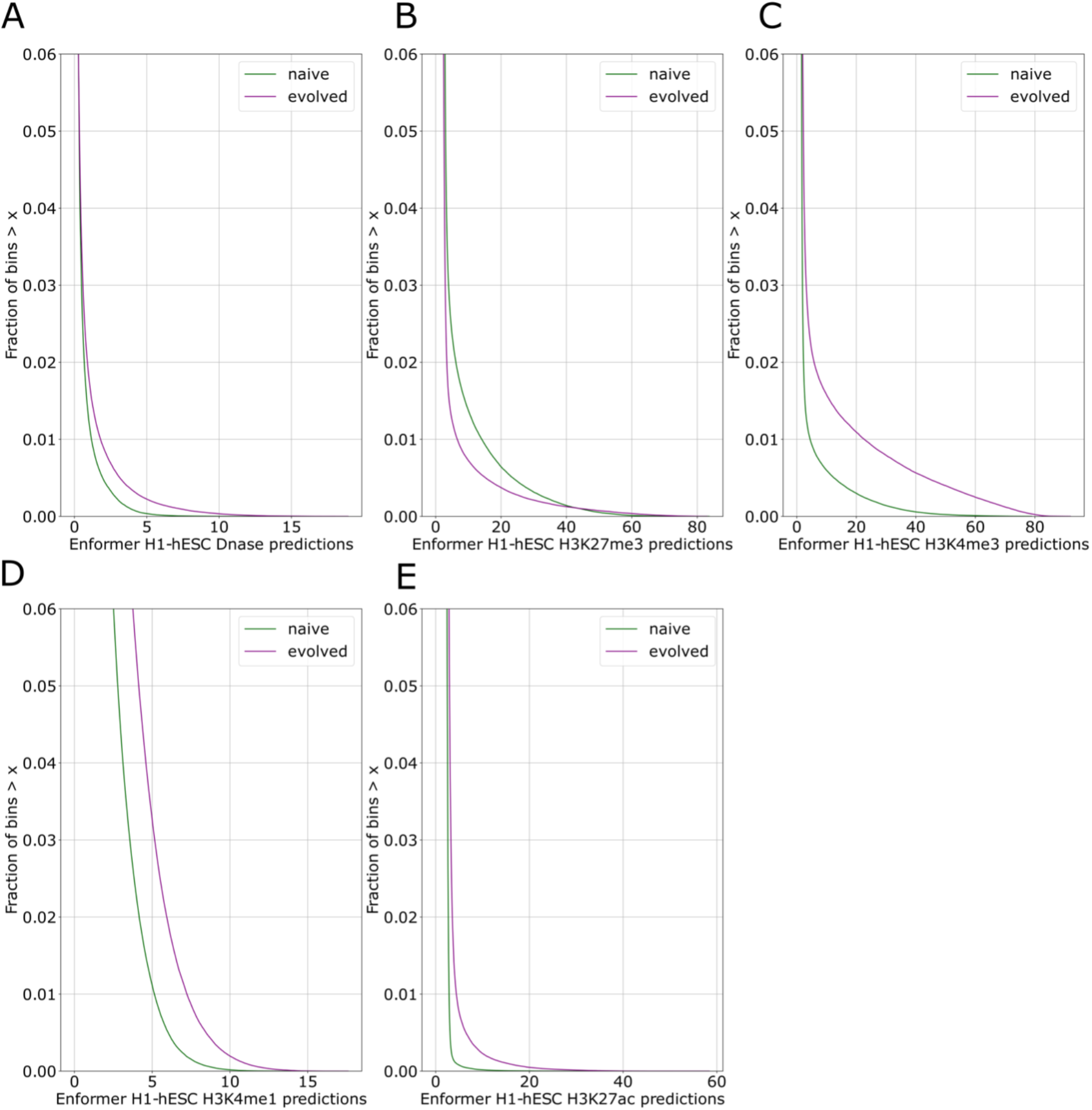
Survival curves for each H1-hESC track. For each predicted Enformer value (*x*-axis), the fraction of bins whose predicted values are at least that large (*y*-axes), for both naïve and evolved (colours). Chromatin marks include (**A**) DNase I, (**B**) H3K27me3, (**C**) H3K4me3, (**D**) H3K4me1, (**E**) H3K27ac.

**Supplementary Figure 13:**
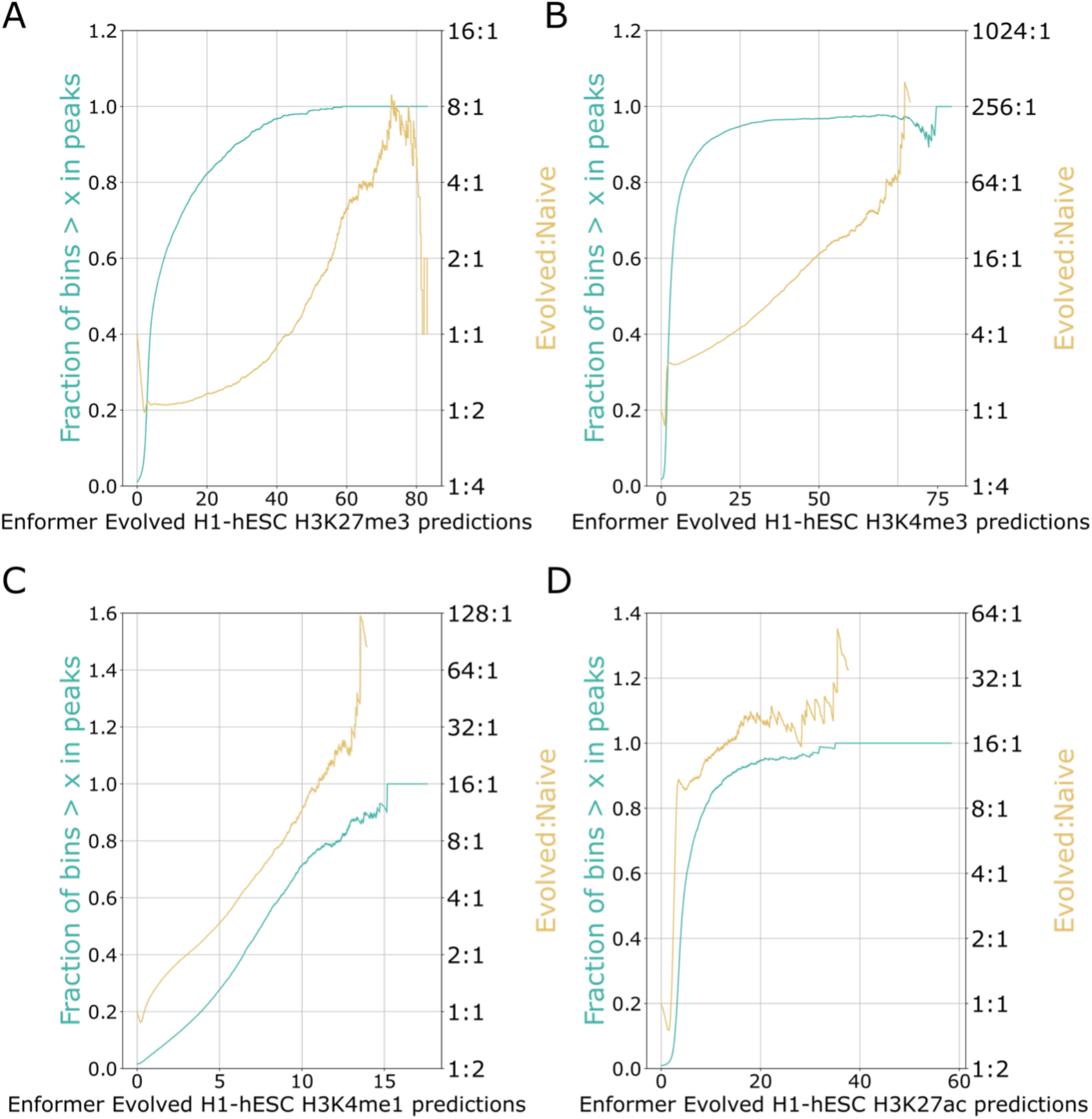
Relative abundances and in peak percentages for each Enformer track. For each Enformer predicted chromatin mark value (*x*-axes), the fraction of prediction bins with at least this value that are in ENCODE peaks (left *y*-axis; cyan) and the ratio of evolved:naïve bins (right *y*-axes; yellow), as in **Fig. 4A**. Chromatin marks include (**A**) H3K27me3, (**B**) H3K4me3, (**C**) H3K4me1, (**D**) H3K27ac.

**Supplementary Table 1:** 12 cell types used in our analysis that had prediction tracks for all 5 Enformer tracks of interest.

| **Cell type** | **Germ Layer** |
| --- | --- |
| iPS_DF_male_newborn | PSC |
| foreskin_melanocyte_male_newborn | Ectoderm |
| neural_progenitor_cell_originated_from_H9 | Ectoderm |
| mesenchymal_stem_cell_originated_from_H1 | Mesoderm |
| iPS_DF_male_newborn_2 | PSC |
| hepatocyte_originated_from_H9 | Endoderm |
| foreskin_keratinocyte_male_newborn | Ectoderm |
| mesendoderm_originated_from_H1 | Prior to germ layer |
| H1-hESC | PSC |
| IMR-90 | Endoderm |
| foreskin_fibroblast_male_newborn | Ectoderm |
| neural_stem_progenitor_cell_originated_from_H1 | Ectoderm |

